# Competition for access to mates predicts female-specific ornamentation and polyandry

**DOI:** 10.1101/2019.12.21.885996

**Authors:** Rosalind L. Murray, Elizabeth J. Herridge, Rob W. Ness, R. Axel W. Wiberg, Luc F. Bussière

## Abstract

Sexually selected ornaments are highly variable and the factors that drive variation in ornament expression are not always clear. Rare instances of female-specific ornament evolution (such as in some dance fly species) are particularly puzzling. While some evidence suggests that such rare instances represent straightforward reversals of sexual selection intensity, the distinct nature of trade-offs between ornaments and offspring pose special constraints in females. To examine whether competition for access to mates generally favours heightened ornament expression, we built a phylogeny and conducted a comparative analysis of Empidinae dance fly taxa that display ornate female-specific ornaments. We show that species with more female-biased operational sex ratios in lek-like mating swarms have greater female ornamentation, and in taxa with more ornate females, polyandry is increased. These findings support the hypothesis that ornament diversity in dance flies depends on female receptivity, which is associated with contests for nutritious nuptial gifts provided by males. Moreover, our results suggest that increases in female receptivity lead to higher levels of polyandry and sperm competition among males. The incidence of both heightened pre-mating sexual selection on females and post-mating selection on males contradicts assertions that sex-roles are straightforwardly reversed in dance flies.

## Introduction

There is a striking diversity of sexual ornaments across animals, even between closely related species (Ord and Stuart-Fox 2006; Pomfret and Knell 2008), but the causes of this diversity are often unclear. An increase in ornamentation is expected to accompany increased sexual selection on the more sexually competitive sex (e.g. Andersson and Iwasa 1996)); for example, if access to receptive mates becomes limited in a species, intense contests for mating opportunities may arise, increasing variation in mating success (Emlen and Oring 1977) and favouring the expression of exaggerated secondary sexual traits that increase attractiveness and mating success (Andersson and Iwasa 1996). What drives increased variation in mating success and whether interspecific variation can fully explain diversity in ornament expression across species is the subject of ongoing debate (e.g. Janicke et al. 2016; Janicke and Morrow 2018).

One critical factor affecting mating system transitions is the intensity of competition for access to mates (i.e. contest intensity). When premating contest intensity is high, variance in mating success increases and can provide strong selection pressures to improve attractiveness in the competing sex. Because access to mates is generally a more strongly limiting factor for the reproductive success of males than it is for females, (Trivers 1972), males tend to be more sexually receptive, which often leads to male biases in the measure of operational sex ratio (OSR; the relative proportion of receptive males and females) (Emlen and Oring 1977; Kvarnemo and Ahnesjo 1996) in most animal species (Janicke et al. 2016). When measures of OSR are male-biased and females are scarce, selection will typically favour traits (such as armaments or ornaments) that improve a male’s access to females. Less is known about how sexual selection operates on females, and whether female-biased sex ratios and increased female competition for males will similarly favour the evolution of attractive female ornaments. In fact, recent work synthesizing studies of many species suggests that while the OSR predicts sexual selection intensity in males, it does not similarly do so for females (Janicke and Morrow 2018).

### Female-specific ornament evolution

Sexual selection in females is a well-documented phenomenon (Clutton-Brock 2009; Shuker 2010; Hare and Simmons 2018). However, female-specific ornament evolution remains uncommon, even among taxa in which females experience relatively strong sexual selection (Amundsen 2000). In the rare cases when female-specific ornaments do evolve (Funk and Tallamy 2000; Charlat et al. 2007; Tobias et al. 2012; Liker et al. 2013; Murray et al. 2017), they are hypothesized to improve attractiveness to mates or as signals in intrasexual competition, just as they do in males (Amundsen 2000). However, there are several theoretical reasons to expect that selection on females need not necessarily directly mirror the situation for males with a male-biased OSR (Gwynne and Simmons 1990; Forsgren et al. 2004; Silva et al. 2010).

Unlike in males, female gamete production is typically very costly (but for a review of variation in spermatogenesis costs see Parker and Pizzari 2010), reproductive success is limited more by gamete production than by access to mates (Trivers 1972), and the expression of expensive ornamental traits could come at a cost to fecundity (Fitzpatrick et al. 1995). Females can overcome any costs associated with ornamentation if they receive direct benefits from mating, such as nutritious nuptial gifts (Vahed 1998; South and Lewis 2012) that compensate for resource investment in ornament expression. However, males might still prefer females who invest less in ornamentation (Fitzpatrick et al. 1995), because by increasing their attractiveness to potential mates, ornamented females are more likely to be polyandrous. Males mating with attractive females are therefore more likely to encounter increased risk or intensity of sperm competition within those females (Herridge et al. 2016). The fact that selection on females might heighten postcopulatory sexual selection on males may help explain why sex differences in sexual selection for female-biased dimorphic traits are not consistently associated with contest intensity (Janicke and Morrow 2018).

### Polyandry and female ornaments

Increased levels of polyandry should produce predictable changes in the reproductive morphology of males, including, for example, increased relative testis investment (Simmons 2001). Relative testis size covaries with rates of female polyandry across diverse taxa (Parker et al. 1997; Vahed et al. 2011). Two complementary hypotheses are frequently invoked to explain this relationship: numerical sperm competition (Gay et al. 2009) and male mating rate (Parker and Ball 2005), (for a review see Vahed and Parker 2011). However, the cause-and-effect relationship between polyandry and testis size may not be as straightforward as originally thought (Simmons 2001); population traits, such as the sex ratio, can influence how a male allocates resources during copulation as well as his relative testis size (Reuter et al. 2008). Therefore, given the complexity of the link between polyandry and relative testis investment, and the rarity of taxa displaying female-specific ornaments, to our knowledge predictions about the relationships between female contest intensity, ornamentation, and male testis investment have never been empirically tested.

In this study, we address these gaps in knowledge by building a molecular phylogeny of some dance fly taxa for which lek-like swarm sex ratios have been measured. We then use a comparative approach to ask 1) how contest intensity (as measured by OSR) covaries with female ornament expression, and 2) how female ornament expression covaries with polyandry (as measured by relative testis investment).

## Methods

### Study Species

Dance flies from the subfamily Empidinae (Diptera: Empididae) include species with almost no sexual dimorphism – typically taxa with male biased mating swarms – as well as species progressively more numerous and elaborate female-specific ornaments in typically female biased mating swarms (Collin 1961; Cumming 1994; Murray et al. 2017). In many dance fly taxa, populations form lek-like mating swarms at specific ‘swarm markers’ throughout the breeding season (Funk and Tallamy 2000, Gwynne; Svensson and Petersson 2000). Male dance flies approach displaying females in the mating swarm (Murray et al. 2018) to offer a nuptial gift (often a prey item) in exchange for copulation. The male passes the nuptial gift to the female prior to copulation, either in flight away from the mating swarm (Funk and Tallamy 2000; Murray et al. 2019) or after landing on vegetation (LeBas and Hockham 2005).

Approximately one third of species from the genera *Rhamphomyia* and *Empis* display female-specific ornamentation (Collin 1961; Cumming 1994). There are three main categories of ornaments: pinnate leg scales, inflatable abdominal sacs and dimorphic wings (in size, colour or both). Pinnate leg scales are the most common form of adornments: hair-like bristles that extend laterally on a female’s legs and are completely absent in male dance flies (Figure 1); they can vary in size and the number of legs on which they occur (e.g. (Collin 1961; Cumming 1994; Gwynne and Bussière 2002; LeBas et al. 2003). Inflatable abdominal sacs are less common (or potentially more difficult to observe), but can also vary considerably in their size and shape (Collin 1961; Cumming 1994; Turner 2012). Sexual dimorphism in wings is reasonably common and highly variable (in both size and colour) across dance fly species. The best-studied examples of exaggerated female wing ornamentation are the enlarged and patterned female wings of *Rhamphomyia marginata* (Miller and Svensson 2014) and the enlarged and darkened wings of *Empis borealis* (Figure 2; Svensson and Petersson 2000).

**Figure 1.**
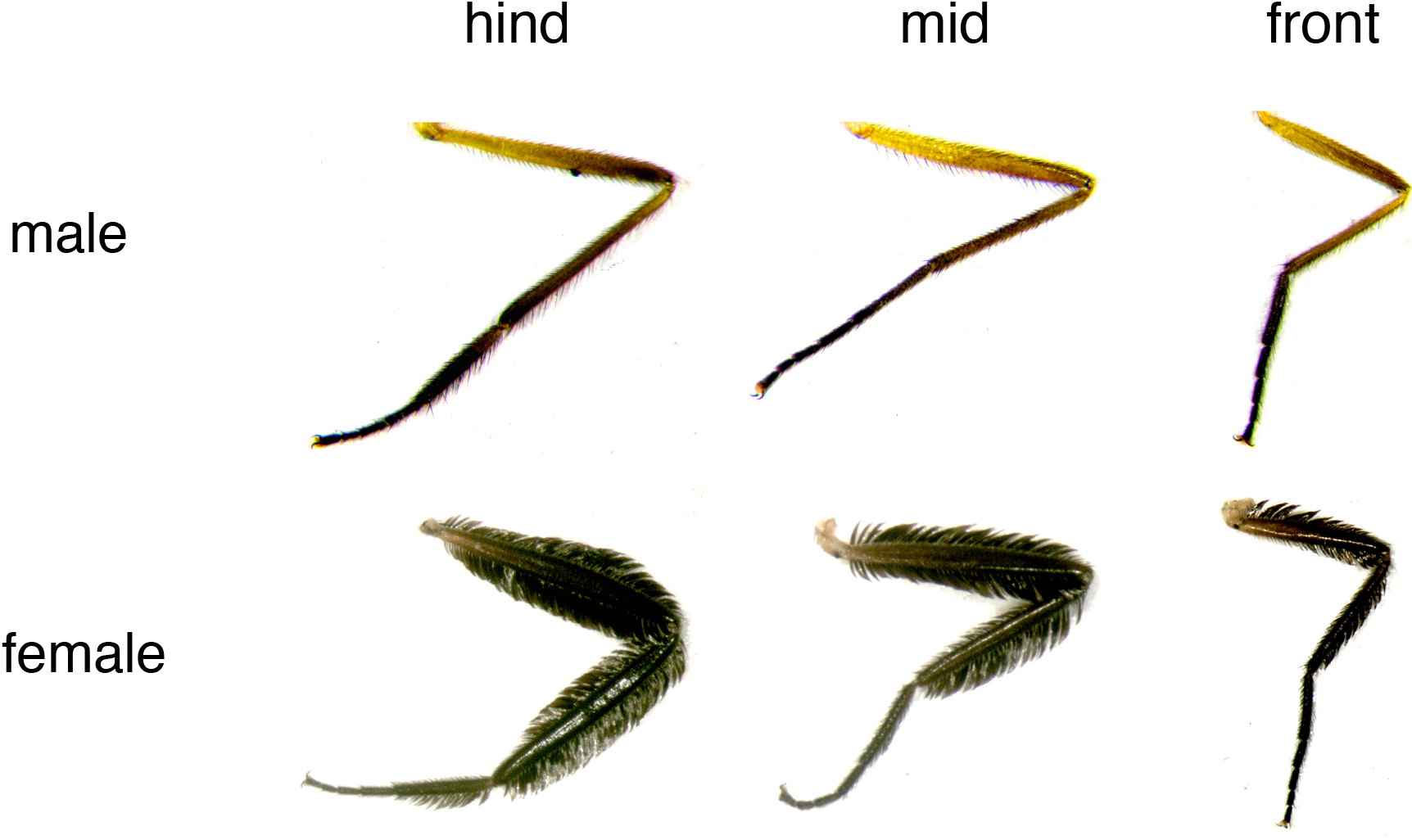
Female-specific pinnate leg scale ornamentation for hind, mid and front legs of *Rhamphomyia longicauda*.

**Figure 2.**
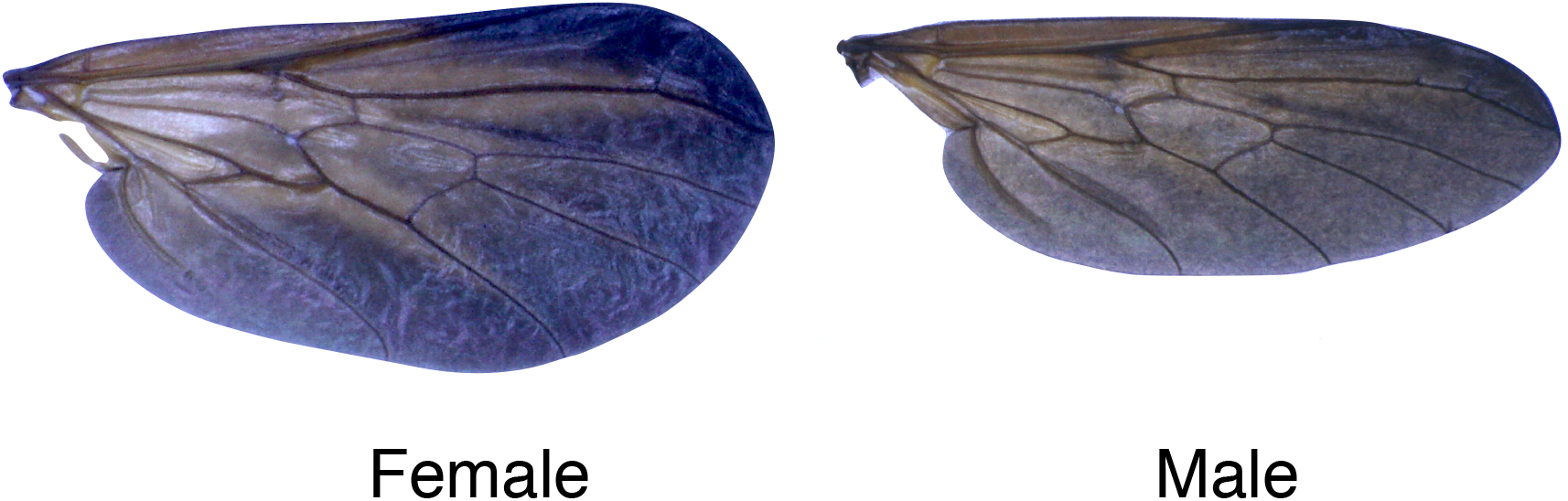
Sexual dimorphism in wing size morphology for *Empis borealis.*

### Sample collections for phylogenetic inference

Although dance flies are common in many parts of the world, swarming behaviour and swarm sex ratios in most species have not been described. We sampled (using sweep nets) 21 species with known swarming sites from Scotland, UK and Ontario, Canada (Table S1). *Empis borealis* samples were collected in the Cairngorms National Park near Aviemore, UK in March 2012 and *Hilara litorea* samples from Edinburgh, UK in July and August 2012. All other UK species were collected from May-July 2011 from the woodland and farmland along the West Highland Way on the eastern side of Loch Lomond between the Scottish Centre for Ecology and the Natural Environment (SCENE) and Rowardennan, Scotland. *Rhamphomyia longicauda* samples were collected in June 2012 from an island in the Credit River near Glen Williams, Ontario, Canada.

### Morphological measurements

Morphological measurements were taken using a dissecting microscope connected to a camera and analyzed using ImageJ (version 1.48) digital imaging software (Abràmoff et al. 2004). To quantify continuous variation in leg and wing dimorphism across species, we took the following external morphological measurements: femora and tibia lengths, wing lengths, thorax lengths, femora and tibia areas, and wing areas. For paired characters we measured both right and left sides and took the mean. When this was not possible because of damage to one side, we measured only the undamaged side.

To estimate sexual dimorphism in pinnate leg scale for each species we took the square root of total leg area (femora and tibia) for each leg (hind, mid and front) and scaled by total leg length. We summed the scaled leg dimorphism values for hind, mid and front legs in each sex. We then subtracted the male value from the female value such that higher positive values of leg dimorphism indicate more pinnation on female legs for that species. Similarly, to quantify wing size dimorphism within each species, we took the square root of wing area and standardised by thorax length for each species. We subtracted the male value from the female value to compute an index of wing size dimorphism for each species.

We constrained the dimorphism measures of legs and wings to positive values because we are interested only in sexual ornament expression (rather than other causes of leg dimorphism) and within these species none of the males are ornamented. Therefore, negative values of leg dimorphism are presumed to be unrelated to ornamentation. The decision to use only positive dimorphism (female-biased) values did not qualitatively affect our analyses, results or interpretations. Finally, we summed continuous measures of leg and wing dimorphism to produce a ‘total ornament’ measure for each species that we used for our ornament measure in the analyses described below.

Wing colour and inflatable abdominal sacs as female ornaments were not measured as continuous traits in this study. We scored wing colour and abdominal sac ornaments as binary (present/absent) traits based on descriptions from the literature (Collin 1961; Cumming 1994). We mapped these discrete traits on to the phylogeny for illustrative purposes, but because of their rarity within our samples and the difficulties in quantifying their expression in the field and across species, we did not include binary ornamentation traits from wing colour or abdominal air sacs in our continuous measures of ‘total ornament’ described above.

To estimate morphological changes reflecting increases in sperm competition that may arise from heightened polyandry, we dissected and measured the two-dimensional area of the testes for ten males per species. We standardized mean testis area by dividing by an individual male’s thorax length (as a proxy for body size) to produce a relative testis size measure for each male. We then took the mean relative testis size of all ten males to use as a species estimate.

### OSR measurements

We estimated the OSR in each species as the number of males swarming divided by the total number of swarming flies. Previous work suggests that mating swarms are often predictable and occur at the same swarm marker for the duration of a swarming season (Funk and Tallamy 2000; Svensson and Petersson 2000), and that the mating swarm OSR can vary both spatially and temporally (Funk and Tallamy 2000; Svensson and Petersson 2000; Wheeler 2008; Murray et al. 2017). We measured multiple swarming events in several species to capture some of the spatial and temporal variation, and to account for sampling error that is especially notable in smaller swarms. In our analyses we computed OSR as a measure of contest intensity in several ways: by summing the total number of swarming OSR tallies for males and females across all swarming events (to derive an “overall swarm sex ratio”), by taking the mean proportion of males across swarming events (i.e., considering each swarm as an independent sample unweighted by swarm size), or by weighting swarms according to the number of swarming flies (to account for the fact that variance is inversely proportional to sample size for binomial samples). Ultimately, we found no qualitative difference in the results regardless of how we summarize our OSR data. The MCMCglmm package in R (Hadfield 2010; R Core Development Team 2014) does not allow for a ‘weights’ argument (as in the lme4 library (Bates et al. 2014)) to account for swarm size. Therefore to account for phylogenetic uncertainty using the MCMCglmm package and the number of animals that were measured to make up each mating swarm sex ratio, below we present analyses that use the summed the total number of swarming OSR tallies across each swarming event for each species.

### Sequencing of CAD for phylogeny estimation

To estimate the evolutionary relationships amongst the 18 fly species of interest for this study, we chose the phylogenetic marker gene CAD (Moulton and Wiegmann 2004). CAD is a fusion protein encoding the first three enzymes of the *de novo* pyrimidine biosynthetic pathway. This gene has proven useful for resolving phylogenetic relationships in Diptera (and particularly in the superfamily Empidoidea) because it is single copy, and possesses moderate levels of non-synonymous divergence (Moulton and Wiegmann 2004, 2007). DNA was isolated from individual flies using DNeasy animal tissue extraction kits (Qiagen, Valencia, CA) according to the manufacturer’s instructions. We amplified a ~1200bp partial coding sequence from the carbomoylphosphate synthase (CPS) domain of CAD using Empididae degenerate PCR primer sequences obtained from personal communication with Brian Cassel from the Wiegmann research group at North Carolina State University (empCAD292F: AGYAATGGNCCNGGHGATCC and empCAD695R: GGRTCYARRTTYTCCATRTTRCA). PCR amplifications were carried out in 20μL reactions with 4.0μL ddH_2_O, 4μL 5X Taq Polymerase Buffer, 1.8-2.1μL of 25 mM MgCl_2_, 2 μL of each primer (2.5μM), 1μL of 10 mM dNTPs, 1 unit of Taq polymerase, and 2-4μL of template DNA. All reactions were carried out using a 3-step touchdown PCR modified from Moulton and Wiegmann (Moulton and Wiegmann 2004): 4 min denaturation at 94°C followed by 4 cycles of 94°C for 30s, 52°C for 30s, 72°C for 2m, 6 cycles of 94°C for 30s, 51°C for 30s, 72°C for 2m, and 36 cycles of 94°C for 30s, 45°C for 20s, 72 °C for 2m30s. PCR amplicons were visualised on 1% agarose gels to ensure that the PCR was successful and generated only a single band. Each PCR fragment was directly sequenced on both strands on an ABI 3730 capillary Sanger sequencing instrument at the Edinburgh Genomics Sequencing facility (Edinburgh, UK). We assembled the forward and reverse strands using Sequencher 4.7 and edited chromatograms manually to ensure that all base calls, and variant sites were reliably scored. We also included the partial CAD sequence of *Hilara lugubris,* which is the only Empidinae species in NCBI Genbank identified to the species level (accession number: DQ369299.1). Our outgroup was *Heterophlebus versabilis* (accession number: HM062728.1) from the related Empididae subfamily, Trichopezinae. This outgroup was chosen because it is closely related to Empidinae based on previous phylogenetic work on the Empidoidea (Moulton and Wiegmann 2007). In addition, the uncorrected pairwise genetic distance between *Heterophlebus versabilis* and each ingroup sequence was always greater than the genetic distance between any pair of ingroup sequences. We aligned all the sequences using their translated amino acid sequences in MUSCLE v 3.8.31 (Edgar 2004) before back converting to a DNA alignment.

### Phylogenetic Inference

We conducted a Bayesian MCMC phylogenetic analysis of CAD sequences in MrBayes v 3.2.4 (Ronquist et al. 2012). Because the CAD sequence we analysed is protein coding, we used a ‘codon’ model of evolution to capture the heterogeneity of mutation rate and selective constraint on sites across the sequence. We also ran simpler models to ensure that we had not over-parameterised the model; we ran one model in which the first, second and third codon positions were partitioned (parameter estimation for each partition was unlinked) and another model where there was no partitioning of sites. For each model we allowed MrBayes to select the best base substitution scheme with a reversible jump MCMC (nst =mixed). We compared the fit of the models by approximating the marginal likelihood with the stepping stone estimator; these marginal-likelihoods were then evaluated using Bayes factors to assess the fit of the data to the three models. For each model we ran three independent runs for 3.5 million cycles, each with four Markov chains. We allowed for a burnin such that the average standard deviation of split frequencies dropped below 0.01 before we began sampling (burnin = 0.5-1.5 million). To ensure convergence, we checked that Potential Scale Reduction Factor (PSRF) for all parameters converged to 1.0 and that the average Estimated Sample Size (ESS) for all parameters exceeded 200. To account for uncertainty in the phylogenetic tree in our statistical analysis, we randomly sampled 1300 topologies from the posterior probability of trees (see below).

For simplicity, we mapped binary ornament traits (see Table 1) onto the consensus tree by delineating the most parsimonious transitions in character states. We note that using binary characters is an oversimplification of the data because even between closely related species that share the same ‘type’ of ornament, there can be substantial variation in ornament expression. In all statistical analyses described below, we use only continuous measures of ornamentation.

**Table 1.** Summary table of morphological traits and operational sex ratio (OSR) across 18 Empidinae dance fly species from three genera (*Empis, Hilara* and *Rhamphomyia*). Continuous traits are displayed as trait mean ± standard error*. OSR is measured as the proportion of males with upper and lower binomial confidence intervals. N=10 for leg, wing and testis measures.

| Species | OSR | Leg dimorphism | Wing dimorphism | Relative testis size | Discrete ornaments** |
| --- | --- | --- | --- | --- | --- |
| *E. aestiva* | 0.34<br>(0.29, 0.39) | 0.178±0.011 | -0.057±0.021 | 0.360±0.018 | PS, WC |
| *E. borealis* | 0.44<br>(0.32, 0.56) | -0.060±0.021 | 0.058±0.008 | 0.147±0.004 | WS |
| *E. nigripes* | 0.46<br>(0.30, 0.62) | 0.094±0.021 | -0.025±0.007 | 0.232±0.015 | PS, WC |
| *E. stercorea* | 0.54<br>(0.23, 0.83) | 0.036±0.031 | -0.0029±0.0008 | 0.225±0.019 | none |
| *E. tessellata* | 0.71<br>(0.61, 0.81) | 0.032±0.008 | 0.025±0.001 | 0.221±0.021 | none |
| *H. beckeri* | 0.75<br>(0.35, 0.96) | 0.0027±0.001 | -0.0041±0.0004 | NA | none |
| *H. chorica* | 0.54<br>(0.52, 0.56) | 0.030±0.011 | 0.017±0.006 | 0.326±0.011 | none |
| *H. litorea* | 0.64<br>(0.62, 0.66) | -0.009±0.007 | 0.011±0.002 | 0.252±0.012 | none |
| *H. interstincta* | 0.82<br>(0.69, 0.91) | -0.0018±0.004 | -0.0024±0.001 | NA | none |
| *H. maura* | 0.62<br>(0.45, 0.79) | -0.050±0.023 | 0.0073±0.0001 | 0.260±0.011 | none |
| *R. crassirostris* | 0.34<br>(0.29, 0.39) | -0.008±0.008 | 0.0096±0.0006 | 0.100±0.010 | none |
| *R. dentipes* | 0.75<br>(0.53, 0.90) | -0.065±0.009 | -0.068±0.009 | 0.195±0.014 | none |
| *R. longicauda* | 0.24<br>(0.20, 0.28) | 0.226±0.008 | 0.0066±0.001 | 0.339±0.007 | PS, AS |
| *R. longipes* | 0.71<br>(0.67, 0.71) | 0.135±0.027 | 0.012±0.008 | 0.253±0.031 | PS |
| *R. nigripennis* | 0.87<br>(0.54, 0.99) | 0.011±0.011 | 0.000064<br>±0.000008 | 0.240±0.022 | WC |
| *R. stigmosa* | 0.57<br>(0.36, 0.78) | 0.021±0.009 | 0.017±0.002 | 0.219±0.006 | none |
| ***R. sulcata*** | 0.63<br>(0.54, 0.99) | -0.035±0.008 | -0.0043±0.0007 | 0.249±0.012 | none |
| ***R. tibiella*** | 0.59<br>(0.46, 0.74) | 0.070±0.004 | -0.014±0.004 | 0.312±0.017 | PS, AS |

\*Continuous traits shown include negative values that were not included in the statistical models where dimorphism measures were constrained to positive values.
\*\*Ornamentation short forms: AS is inflatable abdominal sacs; PS is pinnate leg scales; WS is wing size dimorphism; WC is wing colour dimorphism

### Statistical analyses

All statistical analyses were carried out in R (R Core Development Team 2014). Within each species, all continuous morphological traits were standardized (by subtracting the mean) and scaled (by dividing by two standard deviations) so that each trait was measured on a common scale (Gelman and Hill 2007). The OSR was measured as the logit-transformed ratio (Warton and Hui 2011) of the proportion of males in the mating swarm.

We employed a standard comparative approach (Felsenstein 1985; Harvey and Pagel 1991) to test for an effect of contest intensity (as measured by OSR) on female ornamentation and whether the degree of female ornament expression covaried with male relative testis investment. We used MCMCglmm (Hadfield 2010) to perform comparative analyses using phylogenetic mixed models. We fit two models that initially included fixed effects and their quadratic terms: one with continuous measures of ‘total ornamentation’ as a response (sum of leg and wing size dimorphism described above) and mean-centred OSR and OSR^2^ as fixed effects, and one model that fit testis size as the response predicted by continuous measures of female-specific ornamentation; we did not include OSR measures in our second model (looking for association between polyandry and female ornaments) because our a priori prediction was that ornaments should mediate any relationship between the OSR and testis size, and we wanted to avoid interfering with estimates of that relationship by including a collinear predictor. We fit quadratic terms for sex ratio because contest intensity is predicted to select for ornament expression primarily for the supernumerary sex: as swarms become female biased, we expect ornaments to evolve through female contests for males and their nuptial gifts. However, as the swarm sex ratio becomes male-biased, we predict selection for ornaments to disappear because all females should be able to find mates; the degree to which the swarms are male biased need not covary with ornament expression in this case, since no ornaments are predicted for any male-biased swarm sex ratio. To test for the effect of phylogenetic ancestry, we calculated the phylogenetic heritability (an analogue to Pagel’s lambda), which estimates the proportion of between-species variance explained by the phylogeny (Hadfield 2010).

To correct for uncertainty in the phylogeny during our comparative analysis, we marginalized over the posterior distribution of trees created during phylogenetic inference above. We sampled a tree at iteration t, ran 1000 iterations of the MCMC comparative analysis and then saved the last MCMC sample. The values from the variance components in the saved MCMC sample were then used in the analysis for starting values at iteration t+1 and a new tree from the posterior distribution was taken. This process was repeated 1300 times (i.e. using 1300 trees randomly sampled from the posterior probability of trees) and the first 300 iterations were discarded as burn-in, as in (Ross et al. 2013) while retaining a sample size of 1000.

Our MCMCglmm models assumed a Brownian model on the logit probability scale for the phylogenetic effects (Hadfield 2010). We corrected for phylogenetic non-independence by using the CAD phylogeny as a random effect and the tree sampling method described above. For all models we used a weakly informative parameter-expanded prior. We report the significance of our fixed effects as pMCMC, which is twice the posterior probability that the estimate is positive or negative (whichever is smallest), and can be considered the equivalent of the frequentist p value (Hadfield 2010).

## Results

### Morphological traits and operational sex ratio

We measured the OSR, two continuous measures of female ornamentation (legs and wings), four binary female ornaments and relative testis size across 18 Empidinae dance fly species (Table 1). We found 11 female-specific ornaments among seven dance fly species (the remaining 11 species showed no female ornamentation). The number of mating swarms we sampled per species varied from 2 to 50. Mean OSR measures ranged from very female biased (e.g. *Rhamphomyia longicauda:* 0.24) to very male biased (e.g. *R. longipes:* 0.71). Similarly, standardized leg and wing dimorphism measures ranged from male biased in some species to very female biased (e.g. legs: *R. longicauda;* Figure 1; wings: *E. borealis;* Figure 2). Relative testis size was also variable with a 3-fold increase between the smallest (*R. crassirostris*) to the largest (*E. aestiva*) species measures.

### Empidinae phylogeny

We successfully amplified and sequenced the partial CAD coding sequence for all species included in this study. The chromatograms of *E. stercorea* and *R. stigmosa* were truncated and therefore only partial sequences were included (478bp and 734bp, respectively). All ambiguous bases were marked with an ‘N’ to avoid poor quality nucleotide calls influencing the phylogeny. The alignment of sequences was straightforward with a single 6bp deletion in the ancestor of the *Hilara* species included in our study. We assessed three models of sequence evolution, and found the data fit the ‘codon’ model better (ln(marginal likelihood) = −5988.72) than both simpler models: three codon positions partitioned (ln(marginal likelihood) = −6162.19), and no partitioning (ln(marginal likelihood) = −6516.24). Bayes factor (BF) calculations indicated strong support for the codon model over the partitioned (BF = 2.2 × 10^75^) and unpartitioned (BF = 1.3 × 10^229^) models.

The phylogeny inferred using CAD included 22 species, five *Empis*, six *Hilara*, 10 *Rhamphomyia* and the outgroup *Heterophlebus versabilis*. The outgroup rooted the tree on the branch connecting *Hilara* to the paraphyletic genera *Empis* and *Rhamphomyia*, consistent with Moulton and Wiegmann (2007). The consensus tree displayed in Figure 3 was well resolved: 15 of 19 nodes had a posterior probability >0.95, and only two nodes were unresolved (<0.5), which created a polytomy among *R. crassirostris, R. longicauda* and the well-supported sister pair *R. stigmosa* and *R. sulcata.* Some of these ambiguities could be resolved by including more sequence from CAD and other phylogenetic markers or by sampling more species. However, uncertainty in the exact topology of the unresolved nodes in our CAD tree was accounted for by marginalizing over the posterior probability of tree topologies in our statistical analysis (see below).

**Figure 3.**
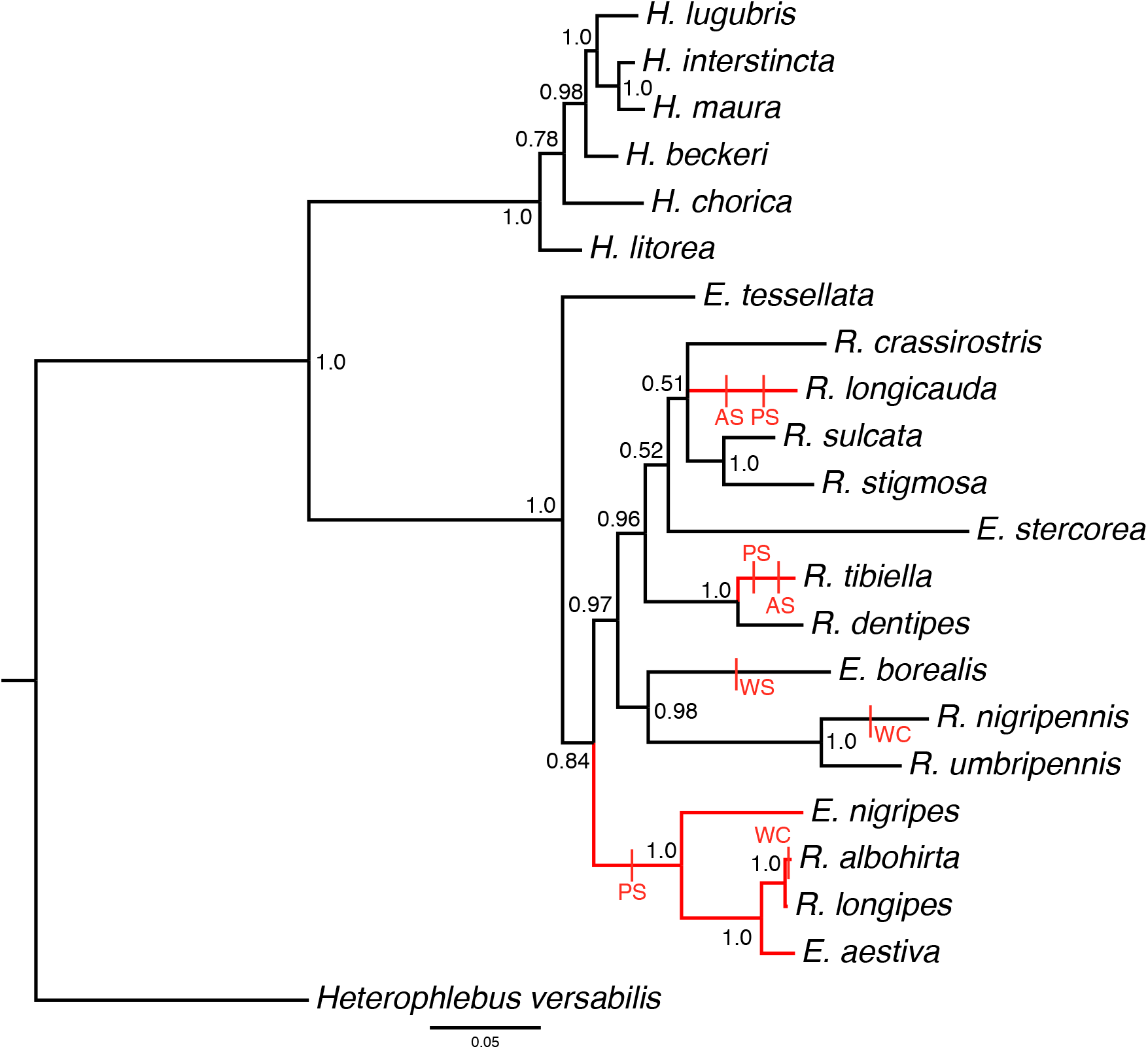
Consensus Bayesian phylogenetic tree of 21 Empidinae species inferred using partial protein coding sequence of CAD. Estimates of the evolutionary relationships between species from the genera *Empis, Hilara,* and *Rhamphomyia* are displayed with the outgroup Heterophlebus versabilis. Node labels indicate the posterior probability of each split, and nodes with less than 0.50 posterior probability are displayed as polytomies. Branches coloured red represent species or clades with female-specific ornamentation in the form of pinnate scales. Red vertical hashes indicate the branch on which different female ornaments are inferred to have arisen. Four ornaments are shown, pinnate scales (PS), inflatable abdominal sacs (AS), wing colour dimorphism (WC) and wing size dimorphism (WS). Each transition in character state was inferred using parsimony.

Our mapping of binary character states, while conservative, does estimate multiple origins of female ornament evolution even within our sample of dance fly species (Figure 3). Pinnate leg scales show three independent origins, wing colour dimorphism and abdominal sacs show two origins, and wing size dimorphism shows a single origin.

### Comparative analysis

If the evolution of female-specific ornaments coevolves with the strength of competition for access to males (or their nuptial gifts), we predicted that increased female ornamentation should positively covary with an increase in female-biased OSR. We fit a phylogenetically controlled generalised linear mixed effects model, with total ornamentation (the sum of wing and leg dimorphism indices obtained for each species) as the response, OSR and OSR^2^ as fixed effects and the phylogeny as a random effect. We found a significant linear (pMCMC=0.018) and quadratic (pMCMC=0.026) association between OSR and female ornamentation after correcting for phylogeny (Table 2): the linear term indicates that as predicted, species with more female-biased swarms had higher levels of ornament expression, while the negative quadratic term indicated a gradual reduction of this effect as the swarm sex ratio approached 0.5. For male biased swarms (OSR >0.5) there was no discernible relationship between the sex ratio and ornament expression (Figure 4). We calculated the phylogenetic heritability (analogue to Pagel’s lambda) as 0.25, indicating a low degree of phylogenetic structure in ornamentation after accounting for variation in OSR measures. The above analysis assumes that the predictors are sampled without error because the MCMCglmm package does not allow for a weighting of the fixed effects (i.e. by number of swarms sampled or sample size within each swarm for our OSR measures). To verify that our estimates were not biased by this assumption, we fit a GLMM in the lme4 package and included number of swarms sampled and separately, number of individuals, per species as ‘weights’ arguments in the models. The results of both models were qualitatively similar (data not shown) to the phylogenetically controlled model.

**Figure 4.**
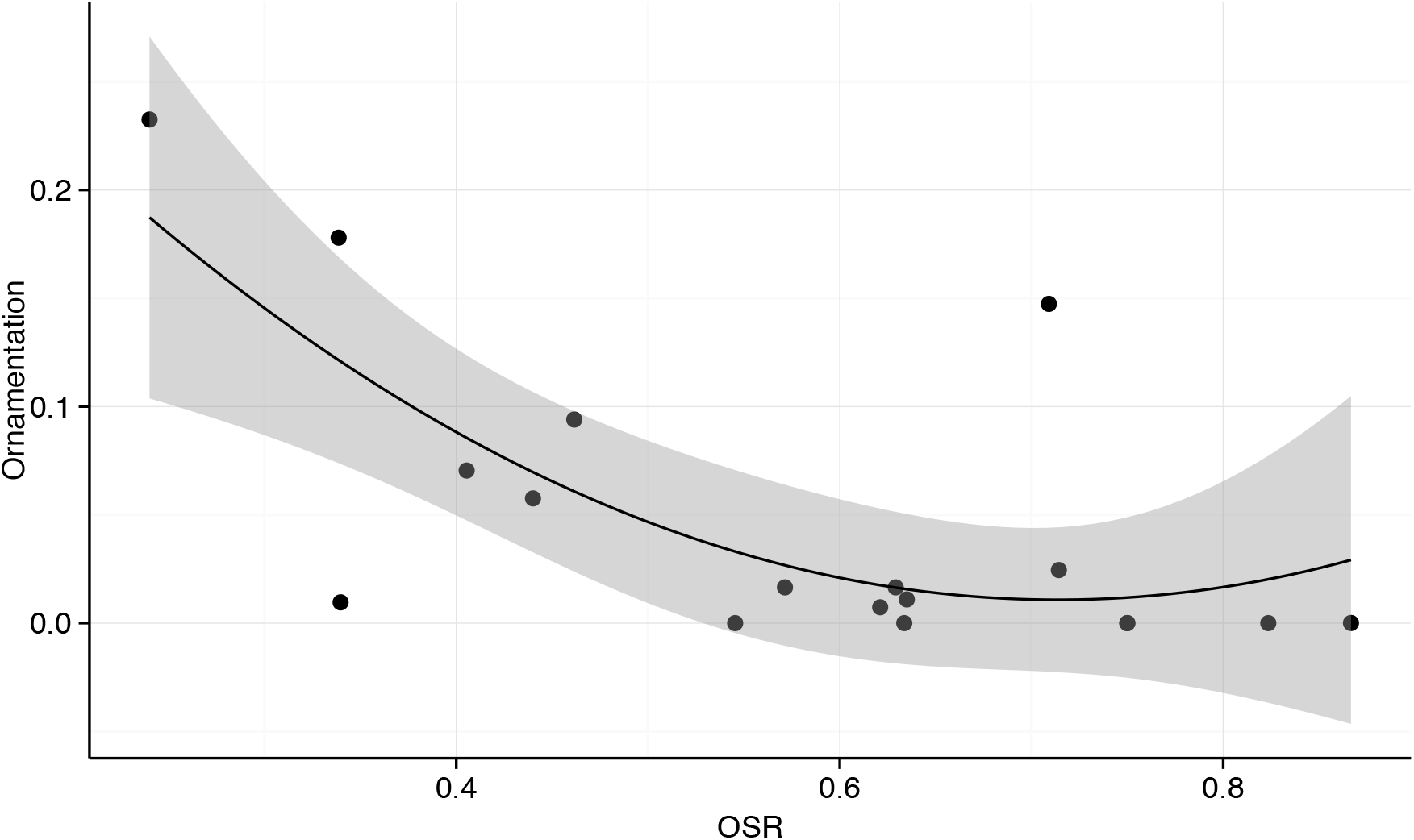
The nonlinear association between competition intensity for mates (measured as operational sex ratio (OSR); proportion of males within a mating swarm) and continuous measures of female-specific ornamentation across dance fly species (see text for details). Lower OSR values indicate female-biased swarms and an OSR of 0.5 indicates equal numbers of males and females. The shaded area is the standard error around the estimate.

**Table 2.** Estimates from a phylogenetically-controlled analysis of female ornamentation (leg dimorphism + wing dimorphism) predicted by operational sex ratio (OSR) estimates across Empidinae species (logit-transformed proportion of males). The model used a Gaussian distribution as specified in MCMCglmm. Values were generated using the summary function in the MCMCglmm package in R.

|  | posterior<br>mean | L-95% CI | U-95% CI | eff. samples | pMCMC |
| --- | --- | --- | --- | --- | --- |
| **Intercept** | 0.038 | -0.017 | 0.089 | 1000 | 0.150 |
| **OSR** | -0.090 | -0.122 | -0.058 | 1000 | <0.001 |
| **OSR^2** | 0.043 | 0.006 | 0.082 | 1000 | 0.022 |

We tested for an association between female ornaments and male testis across dance fly species. We hypothesized that heightened ornament expression to attract males (Murray et al. 2018) might signify a greater level of polyandry, which would in turn select for sperm competition traits in males. We fit a MCMCglmm mixed model with relative testis size as the response, and continuous measures of female leg and wing dimorphism as fixed effects; the quadratic term for ornament expression was removed during model simplification. We also fit phylogeny as a random effect. We found that, as predicted, female ornament expression and relative testis size had a significant positive linear association (pMCMC=0.020). We calculated the posterior probability of phylogenetic signal (mixed model equivalent of Pagel’s lambda; (HADFIELD and NAKAGAWA 2010)) as 0.28, once again indicating a low degree of phylogenetic structure in relative testis size after accounting for variation in ornamentation measures.

## Discussion

Dance flies from the subfamily Empidinae display highly variable female sexual ornaments, ranging from species with multiple female-specific ornaments to those that display very little dimorphism in secondary sex characters (Figure 1, 2). We set out to test two predictions about female ornament evolution: (1) contest intensity (as measured by female-biased OSR) covaries with elaborate female ornament expression, and (2) female ornament expression covaries with polyandry (as measured by relative testis investment). Our phylogeny of 22 species supports multiple origins of female-specific ornaments across species (Figure 3) in line with previous assessments (Cumming 1994; Watts et al. 2016). Comparing taxa in a phylogenetic context we found that (1) pre-mating contest intensity covaried with the degree of female ornamentation and, in line with predictions, only when mating swarms were female-biased (Table 2, Figure 4), and (2) that increased female-specific ornamentation positively covaried with polyandry (inferred from male relative testis investment; Table 3, Figure 5).

**Figure 5.**
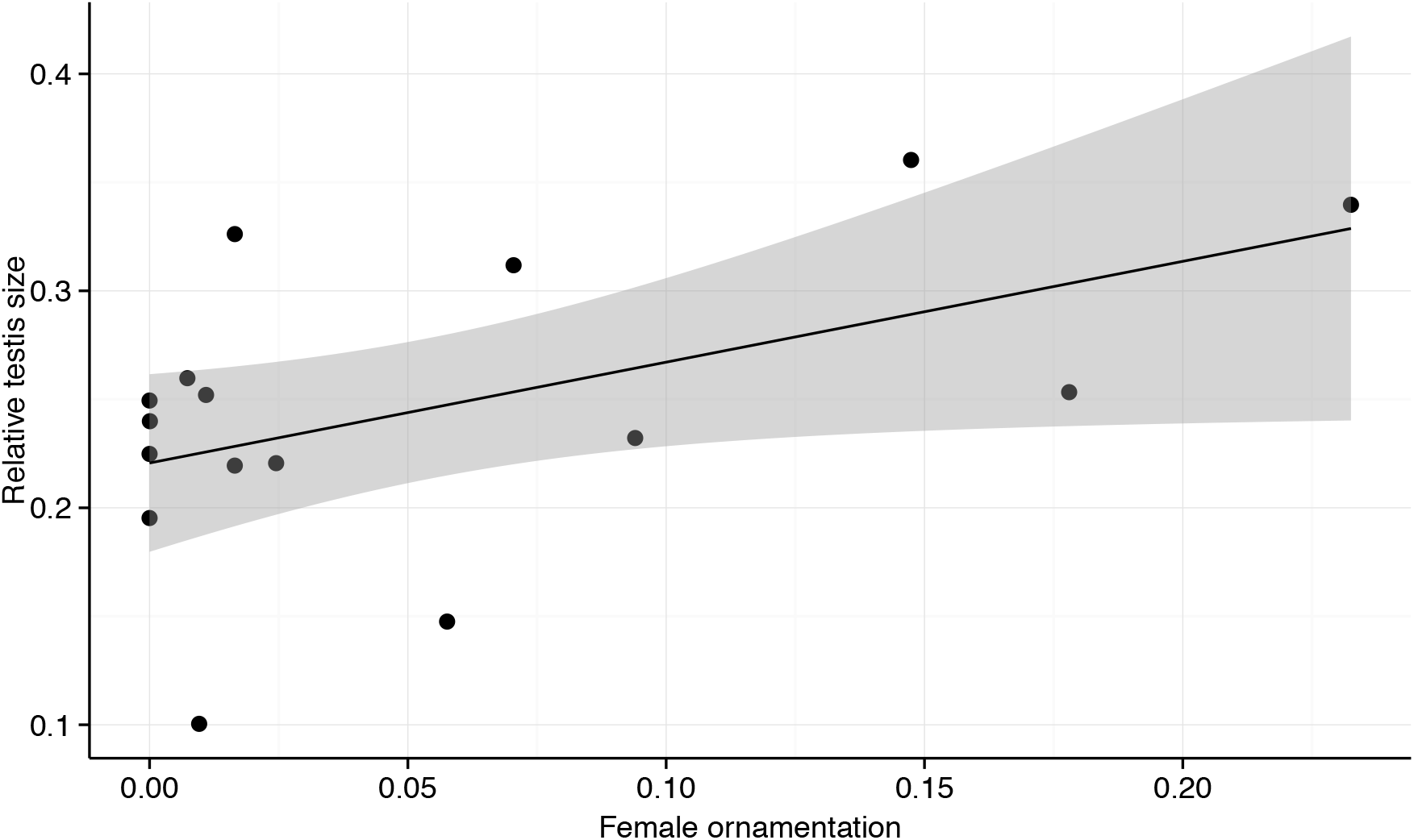
Linear estimate showing the association between female-specific ornamentation and relative testis size across dance fly species (Diptera: Empididae: Empidinae). Shaded area shows the standard error.

**Table 3.** Estimates from a phylogenetically-controlled analysis of female ornaments on the relative testis investment by males across Empidinae species. The model incorporated a “Gaussian” error distribution. Values were generated using the summary function in the MCMCglmm package in R. ‘Ornamentation’ is the female-specific continuous measures of leg and wing dimorphism.

|  | posterior<br>mean | L-95% CI | U-95% CI | eff. samples | pMCMC |
| --- | --- | --- | --- | --- | --- |
| **Intercept** | 0.250 | 0.201 | 0.296 | 1000 | <0.001 |
| **Ornamentation** | 0.032 | 0.006 | 0.062 | 1000 | 0.020 |

### Empidinae phylogeny supports multiple origins of female ornaments

Our phylogeny is consistent with a recent, broad scale tree of the empids (Watts et al. 2016). While the two trees only overlap in 2 species ((Watts et al. 2016) and Figure 3), we do find the same general patterns: all species in the genus *Hilara* are members of a well-supported monophyletic clade, while the genera *Rhamphomyia* and *Empis* are paraphyletic, and form a clade distinct from *Hilara*. We also find that there are multiple independent transitions, both from unornamented ancestors to extant taxa with one or multiple female-specific ornaments, and within each ornament class (pinnate leg scales, abdominal sacs and wing size and colour dimorphism; Table 1, Figure 3) It is important to note that this tree represents only a small fraction of the species diversity in the Empidinae (Watts et al. 2016) and more complete taxonomic sampling could alter the mapping of these traits and our inferences about how many times ornamentation has evolved (see review by Nabhan and Sarkar 2012). However, given that we report four distinct sexually dimorphic traits that all appear to have arisen independently, it seems likely that female-specific ornamentation has arisen multiple times within the dance flies. In addition, the pinnate scales of *R. longicauda* and *R. tibiella,* which appear as separate transitions, are morphologically distinct and unlikely to be homologous traits (Collin 1961); Figure 1). Indeed, even for *R. longipes and R. albohirta,* which our tree identifies as sister species that share the same origin of leg pinnation (Figure 3), the degree of ornamentation is different; *R. longipes* displays hind leg pinnation, while *R. albohirta* females have pinnation on their hind and mid legs (Collin 1961). Therefore, by simplifying continuous ornamental traits into binary characters on the phylogeny we are being conservative in estimating the extent of ornament evolution in the dance flies. Indeed, given the variation in traits coded as the same in Table 1 and Figure 3, it is likely that each class or ornament is quite labile through evolutionary time. A complete comparative analysis with more thorough phenotypic sampling of the *Empis* and *Rhamphomyia* clade would provide valuable insights into the evolution of different forms of female-specific ornamentation.

### Female-biased contest intensity and female ornamentation

For the first time, we combine phylogenetic analyses with observations of swarm attendance by males and females to estimate female contest intensity and its role in driving ornament expression. We found a significant nonlinear relationship between the amount of female-female competition for access to mates (as measured by the OSR) and the amount of female-specific ornamentation (Table 2, Figure 4). Our results show that ecological processes observable at the species level (contest intensity in mating swarms) reflect patterns at the macroevolutionary scale: on average, species with more intense intrasexual contests for mates possess more exaggerated female-specific ornaments. Furthermore, we document that this relationship appears to be constrained to female-biased swarms: the relationship between female contest intensity and ornament expression dissipates as the sex ratio approaches 0.5. This finding is notable not least because recent work by Janicke and Morrow (2018) found no evidence for nonlinearity in a deeper macroevolutionary analysis of how sex ratio covaries with the intensity of sexual selection; clearly the prediction of nonlinearity can hold within some groups even if it is absent across a wider survey of animals.

Our finding that female-biased OSR measures associate with female ornaments is qualitatively similar to a study by Pomfret and Knell (2008) in male *Onthophagus* beetles; they observed that an increase in intrasexual competition was associated with the presence of male weaponry. Our study supports the idea that sexual selection on females can result in similar outcomes to those observed in males (reviewed by Hare and Simmons 2018); females evolve ornaments when they are involved in intense intrasexual competition. Interestingly, a recent meta-analysis by Janicke and Morrow (2018) suggested that the OSR is a useful predictor of the opportunity for sexual selection in males, but not females. Our findings differ from the meta-analysis results further because we found a low phylogenetic signal (0.25) for the relationship between OSR and ornamentation, while their study found a strong phylogenetic signal (Pagel’s lambda: 0.42-0.95 depending on the metric of sexual selection measured (Janicke and Morrow 2018)). This difference in phylogenetic signal between studies might be related to the evolutionary timescale of these studies (i.e. Empidinae subfamily compared to Animal kingdom), and/or due to the variation of dance fly mating behaviour that makes these taxa more labile in female ornamentation.

### Operational sex ratio as a measure of contest intensity in dance flies

As measures of the OSR become increasingly biased toward one sex, the intensity of both intra- and intersexual selection on that sex is expected to rise (Emlen and Oring 1977; Gwynne 1990; Clutton-Brock and Parker 1992; Johnstone et al. 1996; Kvarnemo and Ahnesjo 1996, 2002), and several studies have shown that the OSR as a measure can accurately reflect levels of intrasexual competition (Jirotkul 1999; Berglund and Rosenqvist 2008; Silva et al. 2010; Monteiro and Lyons 2012; Monteiro and Vieira 2013; Janicke and Morrow 2018). One assumption for the OSR to reliably indicate the intensity of sexual selection is that an increase in the bias of the OSR increases mate monopolization by the more common sex (Emlen and Oring 1977). However, mate monopolization, while potentially very important for male reproductive success (but see Janicke and Morrow 2018), is unlikely to be tightly linked to female reproductive success. Instead, females are more likely to be limited by access to resources than sperm (Trivers 1972). Male dance flies provide nutritious nuptial gifts to their mates at copulation (Collin 1961; Cumming 1994). Recently, Hunter and Bussière (Hunter and Bussiere 2018) showed that females of an ornamented dance fly species could only mature their eggs if they mated (and received a nuptial gift), while this was not the case for an unornamented species. Therefore, the sometimes-intense contests for mates we observe among females (Table 1, Figure 4) are very likely associated with selection to obtain nutritious nuptial gifts to promote ovarian maturation (Hunter and Bussiere 2018). The skewed swarm sex ratios that arise due to these contests therefore reliably predict the degree of female-specific ornamentation in most (but not all) species; e.g. *R. longipes* appears as a clear outlier in our dataset as an ornamented species with a strongly-male biased sex ratio (Table 1), that potentially represents a recent transition in mating system.

### Female-specific ornamentation and male relative testis investment

Our study supports the idea that in dance flies there is a strong relationship between premating sexual selection on females and post-mating sexual selection on males. We show that taxa where females with more intense levels of contests for access to mates are more likely to evolve attractive ornaments and mate with multiple males. We found a positive association between female-specific ornamentation and male relative testis size across species (Table 3, Figure 5). One of the most consistently observed patterns relating to increased testis size is a positive covariance with polyandry; to be more specific, testis size increases when females mate with more than one male (Pitnick 1996; Pitcher et al. 2005; Montgomerie and Fitzpatrick 2009; Soulsbury 2010; Vahed et al. 2011). Two hypotheses (reviewed in Vahed and Parker (2011), that may be interrelated, can account for the testis size-polyandry covariance. The numerical sperm competition hypothesis posits that larger testes will allow males to produce more sperm per ejaculate thus competing more effectively (Parker et al. 1997), while the male mating rate hypothesis predicts that males with larger testes will be able to increase the number of copulations they engage in. Further studies examining the relationship between ejaculate investment, male mating rate and female ornamentation would be necessary to identify which hypothesis is more likely to explain the relationship observed in the dance flies. Critically, male dance flies are likely limited in how often they can mate by how quickly they are able to acquire a new prey-item nuptial gift (Cumming 1994; LeBas et al. 2014) and return to the mating swarm (See (Kokko et al. 2012)). The ‘time away’ from the mating swarm that males require to collect nuptial gifts could limit the likelihood that male mating rate is contributing to the increased relative testis size observed in ornamented species of dance flies.

The existence of the relationship between increased female ornaments and increased relative testis investment in dance flies is additionally notable because of the uncertainty in how ornament expression covaries with female mating frequency. While female ornaments are known to improve attractiveness in some species (Funk and Tallamy 2000; LeBas et al. 2003; Murray et al. 2018), in other cases, the relationship between female ornament size and female probability of mating (Wheeler et al. 2012) or number of mates (Herridge 2016) is not as clear. One possible explanation for the discrepancy between attractiveness and mating success in dance flies is coevolutionary sexual conflict; ornamented females are eager to accept nuptial gifts, while discriminating males are reluctant to mate with highly polyandrous females because they pose an increased risk of sperm competition. Our results suggest that female ornaments evolve in species with the highest mating rates whatever the current status of coevolution between the sexes in any particular species.

Our work shows that female contest intensity covaries with female ornament expression and that females with larger ornaments are likely polyandrous. Within the dance flies these elaborate female traits appear to be highly labile; multiple evolutionary origins of variable sizes and types of ornaments. Importantly, OSR predicts ornament expression even in systems with female ornaments, but there might not be an overall sex difference in sexual selection because the very process that favours ornaments in females also promotes postcopulatory sexual selection in males. This key difference may help explain otherwise perplexing patterns that emerge from comparisons across the animal kingdom

## Acknowledgements

We would like to thank D. Gwynne for his insightful contributions and support during the development of this work. We also thank B. Cassel, T. Houslay, P. Keightley, P. Lee, T. Little, A. Plant, K. Vahed and B. Wiegmann for their assistance and the Scottish Centre for Ecology and the Natural Environment (SCENE) and the Gwynne field stations. This work was supported by the Royal Society of London, NSERC, the University of Stirling and the University of Toronto.

## Supplementary

**Table S1.** Sample collection locations for Empidinae dance flies.

| Species | GPS coordinates | Collection locations |
| --- | --- | --- |
| *Empis aestiva* | 56°09'06.35"N, 004°38'36.20"W | SCENE, UK |
| *Empis borealis* | 57°14'38.20"N, 003°41'42.55"W | Aviemore, UK |
| *Empis nigripes* | 56°09'06.35"N, 004°38'36.20"W | SCENE, UK |
| *Empis stercorea* | 56°09'06.35"N, 004°38'36.20"W | SCENE, UK |
| *Empis tessellata* | 56°09'06.35"N, 004°38'36.20"W | SCENE, UK |
| *Hilara chorica* | 56°09'06.35"N, 004°38'36.20"W | SCENE, UK |
| *Hilara litorea* | 55°55'22.28"N, 003°10'34.04"W | Edinburgh, UK |
| *Hilara maura* | 56°09'06.35"N, 004°38'36.20"W | SCENE, UK |
| *Rhamphomyia crassirostris* | 56°09'06.35"N, 004°38'36.20"W | SCENE, UK |
| *Rhamphomyia dentipes* | 56°09'06.35"N, 004°38'36.20"W | SCENE, UK |
| *Rhamphomyia longicauda* | 43°41'11.00"N, 079°55'34.00"W | Glen Williams, Canada |
| *Rhamphomyia longipes* | 56°09'06.35"N, 004°38'36.20"W | SCENE, UK |
| *Rhamphomyia nigripennis* | 56°09'06.35"N, 004°38'36.20"W | SCENE, UK |
| *Rhamphomyia stigmosa* | 56°09'06.35"N, 004°38'36.20"W | SCENE, UK |
| *Rhamphomyia sulcata* | 56°09'06.35"N, 004°38'36.20"W | SCENE, UK |
| *Rhamphomyia tibiella* | 56°09'06.35"N, 004°38'36.20"W | SCENE, UK |

## References

Abràmoff, M. D., P. J. Magalhães, and S. J. Ram. 2004. Image processing with ImageJ. Biophotonics international 11:36–43.

Amundsen, T. 2000. Why are female birds ornamented? Trends in Ecology and í Evolution 15:149–155.

Andersson, M. and Y. Iwasa. 1996. Sexual selection. Trends Ecol. Evol. 11:53–58.

Bates, D., M. Mächler, B. Bolker, and S. Walker. 2014. Fitting linear mixed-effects models using lme4. arXiv 1406.5823.

Berglund, A. and G. Rosenqvist. 2008. An intimidating ornament in a female ! pipefish. Behavioral Ecology 20:54–59.

Charlat, S., M. Reuter, E. A. Dyson, E. A. Hornett, A. Duplouy, N. Davies, G. K. Roderick, N. Wedell, and G. D. D. Hurst. 2007. Male-killing bacteria trigger i a cycle of increasing male fatigue and female promiscuity. Current Biology 17:273–277.

Clutton-Brock, T. 2009. Sexual selection in females. Animal Behaviour 77:3–11.

Clutton-Brock, T. H. and G. A. Parker. 1992. Potential reproductive rates and the operation of sexual selection. Quarterly Review of Biology:437–456.

Collin, J. E. 1961. British Flies VI: Empididae Part 2: Hybotinae, Empidinae (except Hilara). Cambridge University Press, Cambridge, UK.

Cumming, J. M. 1994. Sexual selection and the evolution of dance fly mating systems (Diptera: Empididae; Empidinae). The Canadian Entomologist 126:907–920.

Edgar, R. C. 2004. MUSCLE: a multiple sequence alignment method with í reduced time and space complexity. BMC Bioinformatics 5:113.

Emlen, S. T. and L. W. Oring. 1977. Ecology, sexual selection, and the evolution of mating systems. Science 197:215–223.

Felsenstein, J. 1985. Phylogenies and the comparative method. Am. Nat. 125:1–15.

Fitzpatrick, S., A. Berglund, and G. Rosenqvist. 1995. Ornaments or offspring: costs to reproductive success restrict sexual selection processes. Biological Journal of the Linnean Society 55:251–260.

Forsgren, E., T. Amundsen, Å. A. Borg, and J. Bjelvenmark. 2004. Unusually dynamic sex roles in a fish. Nature 429:551–554.

Funk, D. H. and D. W. Tallamy. 2000. Courtship role reversal and deceptive signals in the long-tailed dance fly, Rhamphomyia longicauda. Animal Behaviour 59:411–421.

Gay, L., D. J. Hosken, R. Vasudev, T. Tregenza, and P. E. Eady. 2009. Sperm competition and maternal effects differentially influence testis and sperm size in Callosobruchus maculatus. Journal of Evolutionary Biology 22:1143–1150.

Gelman, A. and J. Hill. 2007. Replication data for: data analysis using regression and multilevel/hierarchical models. Institute for Quantitative Social Science.

Gwynne, D. T. 1990. Testing parental investment and the control of sexual selection in katydids: the operational sex ratio. Am. Nat. 136:474–484.

Gwynne, D. T. and L. T. Bussière. 2002. Female mating swarms increase predation risk in a role-reversed dance fly (Diptera: Empididae: *Rhamphomyia longicauda Loew)*. Behaviour 139:1425–1430.

Gwynne, D. T. and L. W. Simmons. 1990. Experimental reversal of courtship roles in an insect. Nature 346:172–174.

Hadfield, J. D. 2010. MCMC methods for multi-response generalized linear mixed models: the MCMCglmm R package. Journal of Statistical Software 33:1–22.

Hadfield, J. D. and S. Nakagawa. 2010. General quantitative genetic methods for comparative biology: phylogenies, taxonomies and multi-trait models for continuous and categorical characters. Journal of Evolutionary Biology 23:494–508.

Hare, R. M. and L. W. Simmons. 2018. Sexual selection and its evolutionary consequences in female animals. Biological Reviews:000-000.

Harvey, P. H. and M. D. Pagel. 1991. The comparative method in evolutionary biology. Oxford Press, Oxford, UK.

Herridge, E. J. 2016. The role of polyandry in sexual selection among dance flies. Biological and Environmental Sciences. University of Stirling, Stirling, UK.

Herridge, E. J., R. L. Murray, D. T. Gwynne, and L. F. Bussiere. 2016. Diversity in mating and parental sex roles. Encyclopedia of Evolutionary Biology:453–458.

Hunter, F. D. L. and L. F. Bussiere. 2018. Comparative evidence supports a role for reproductive allocation in the evolution of female ornament diversity. Ecol. Entomol.

Janicke, T., I. K. Haderer, M. J. Lajeunesse, and N. Anthes. 2016. Darwinian sex roles confirmed across the animal kingdom. Science Advances 2.

Janicke, T. and E. H. Morrow. 2018. Operational sex ratio predicts the opportunity and direction of sexual selection across animals. Ecology Letters 21:384–391.

Jirotkul, M. 1999. Operational sex ratio influences female preference and male-male competition in guppies. Animal Behaviour 58:287–294.

Johnstone, R. A., J. D. Reynolds, and J. C. Deutsch. 1996. Mutual mate choice and sex differences in choosiness. Evolution 50:1382–1391.

Kokko, H., H. Klug, and M. D. Jennions. 2012. Unifying cornerstones of sexual selection: operational sex ratio, Bateman gradient and the scope for competitive investment. Ecology Letters 15:1340–1351.

Kvarnemo, C. and I. Ahnesjo. 1996. The dynamics of operational sex ratios and competition for mates. Trends Ecol. Evol. 11:404–408.

Kvarnemo, C. and I. Ahnesjo. 2002. Operational sex ratios and mating competition. Sex Ratios: Concepts and Research Methods:366–382.

LeBas, N. R. and L. R. Hockham. 2005. An Invasion of cheats. Current Biology 15:64–67.

LeBas, N. R., L. R. Hockham, and M. G. Ritchie. 2003. Nonlinear and correlational sexual selection on honest female ornamentation. Proceedings of the Royal Society B: Biological Sciences 270:2159–2165.

LeBas, N. R., L. R. Hockham, and M. G. Ritchie. 2014. Sexual selection in the gift-giving dance fly, Rhamphomyia sulcata, favors small males carrying small gifts. Evolution 58:1763–1772.

Liker, A., R. P. Freckleton, and T. Székely. 2013. The evolution of sex roles in birds is related to adult sex ratio. Nature Communications 4:1587.

Miller, C. W. and E. I. Svensson. 2014. Sexual Selection in Complex Environments. Annual Review of Entomology 59:427–445.

Monteiro, N. M. and D. O. Lyons. 2012. Stronger sexual selection in warmer waters: the case of a sex role reversed pipefish. PLoS One 7:9.

Monteiro, N. M. and M. N. Vieira. 2013. Operational sex ratio, reproductive costs, and the potential for intrasexual competition. Biological Journal of the Linnean Society 110:477–484.

Montgomerie, R. and J. L. Fitzpatrick. 2009. Testes, sperm, and sperm competition. Science Publishers.

Moulton, J. K. and B. M. Wiegmann. 2004. Evolution and phylogenetic utility of CAD (rudimentary) among Mesozoic-aged Eremoneuran Diptera (Insecta). Molecular Phylogenetics and Evolution 31:363–378.

Moulton, J. K. and B. M. Wiegmann. 2007. The phylogenetic relationships of flies in the superfamily Empidoidea (Insecta: Diptera). Molecular Phylogenetics and Evolution 43:701–713.

Murray, R. L., D. T. Gwynne, and L. F. Bussiere. 2019. The role of functional constraints in nonrandom mating patterns for a dance fly with female ornaments. Journal of Evolutionary Biology 32:984–993.

Murray, R. L., E. J. Herridge, R. W. Ness, and L. F. Bussiere. 2017. Are sex ratio distorting endosymbionts responsible for mating system variation among dance flies (Diptera: Empidinae)? PLoS One 12:e0178364.

Murray, R. L., J. Wheeler, D. T. Gwynne, and L. F. Bussiere. 2018. Sexual selection on multiple ornaments in dance flies. Proc. R. Soc. B-Biol. Sci. 285:20181525.

Nabhan, A. R. and I. N. Sarkar. 2012. The impact of taxon sampling on phylogenetic inference: a review of two decades of controversy. Briefings in Bioinformatics 13:122–134.

Ord, T. J. and D. Stuart-Fox. 2006. Ornament evolution in dragon lizards: multiple gains and widespread losses reveal a complex history of evolutionary change. Journal of Evolutionary Biology 19:797–808.

Parker, G. A. and M. A. Ball. 2005. Sperm competition, mating rate and the evolution of testis and ejaculate sizes: a population model. Biology Letters 1:235–238.

Parker, G. A., M. A. Ball, P. Stockley, and M. J. G. Gage. 1997. Sperm competition games: a prospective analysis of risk assessment. Proc. R. Soc. B-Biol. Sci. 264:1793–1802.

Parker, G. A. and T. Pizzari. 2010. Sperm competition and ejaculate economics. Biological Reviews 85:897–934.

Pitcher, T. E., P. O. Dunn, and L. A. Whittingham. 2005. Sperm competition and the evolution of testes size in birds. Journal of Evolutionary Biology 18:557–567.

Pitnick, S. 1996. Investment in testes and the cost of making long sperm in Drosophila. Am. Nat. 148:57–80.

Pomfret, J. C. and R. J. Knell. 2008. Crowding, sex ratio and horn evolution in a South African beetle community. Proceedings of the Royal Society B: Biological Sciences 275:315–321.

R Core Development Team. 2014. R: A Language and Environment for Statistical Computing. R Foundation for Statistical Computing, Vienna, Austria.

Reuter, M., J. R. Linklater, L. Lehmann, K. Fowler, T. Chapman, and G. D. D. Hurst. 2008. Adaptation to experimental alterations of the operational sex ratio in populations of Drosophila melanogaster. Evolution 62:401–412.

Ronquist, F., M. Teslenko, P. van der Mark, D. L. Ayres, A. Darling, S. Hohna, B. Larget, L. Liu, M. A. Suchard, and J. P. Huelsenbeck. 2012. MrBayes 3.2: Efficient Bayesian phylogenetic inference and model choice across a large model space. Syst. Biol. 61:539–542.

Ross, L., A. Gardner, N. Hardy, and S. A. West. 2013. Ecology, not the genetics of sex determination, determines who helps in eusocial populations. Current Biology 23:2383–2387.

Shuker, D. M. 2010. Sexual selection: endless forms or tangled bank? Pp. e11–e17. Animal Behaviour. Elsevier Ltd.

Silva, K., M. N. Vieira, V. C. Almada, and N. M. Monteiro. 2010. Reversing sex role reversal: compete only when you must. Animal Behaviour 79:885–893.

Simmons, L. W. 2001. Sperm competition and its evolutionary consequences in the insects. Princeton University Press, Princeton, New Jersey, USA.

Soulsbury, C. D. 2010. Genetic patterns of paternity and testes size in mammals. PLoS One 5: e9581.

South, A. and S. M. Lewis. 2012. Determinants of reproductive success across sequential episodes of sexual selection in a firefly. Proc. R. Soc. B-Biol. Sci. 279:3201–3208.

Svensson, B. G. and E. Petersson. 2000. Swarm site fidelity in the sex role-reversed dance fly Empis borealis. Journal of Insect Behavior 13:785–796.

Tobias, J. A., R. Montgomerie, and B. E. Lyon. 2012. The evolution of female ornaments and weaponry: social selection, sexual selection and ecological competition. Philosophical Transactions of the Royal Society B: Biological Sciences 367:2274–2293.

Trivers, R. L. 1972. Mother-offspring conflict. Am. Zool. 12:648–648.

Turner, S. P. 2012. The Evolution of Sexually Selected Traits in Dance Flies. Pp. 87. Entomology. North Carolina State University, Raleigh, North Carolina.

Vahed, K. 1998. The function of nuptial feeding in insects: review of empirical studies. Biological Reviews of the Cambridge Philosophical Society 73:43–78.

Vahed, K. and D. J. Parker. 2011. The evolution of large testes: sperm competition or male mating rate? Ethology 118:107–117.

Vahed, K., D. J. Parker, and J. D. J. Gilbert. 2011. Larger testes are associated with a higher level of polyandry, but a smaller ejaculate volume, across bushcricket species (Tettigoniidae). Biology Letters 7:261–264.

Warton, D. I. and F. K. C. Hui. 2011. The arcsine is asinine: the analysis of proportions in ecology. Ecology 92:3–10.

Watts, M., I. S. Winkler, C. Daugeron, C. J. B. de Carvalho, S. P. Turner, and B. M. Wiegmann. 2016. Where do the Neotropical Empidini lineages (Diptera: Empididae: Empidinae) fit in a worldwide context? Pp. 67–78. Molecular Phylogenetics and Evolution.

Wheeler, J. 2008. Sexual selection and female ornamentation in a role-reversed dance fly. Pp. 87. Ecology and Evolutionary Biology. University of Toronto.

Wheeler, J., D. T. Gwynne, and L. F. Bussière. 2012. Stabilizing sexual selection for female ornaments in a dance fly. Journal of Evolutionary Biology 25:1233–1242.

